# The *in vitro* bacterial viability and microbial composition of commercially available canine and feline fecal microbial transplantation products

**DOI:** 10.1101/2025.11.10.687729

**Authors:** Nina K. Randolph, Lisa Wetzel, Dubraska Diaz-Campos, Joany C. van Balen, Jenessa A. Winston

## Abstract

Fecal microbial transplantation (FMT) is the transfer of feces from a healthy donor into the gastrointestinal tract of a diseased recipient to confer a health benefit. FMT is increasingly utilized in veterinary medicine and is offered commercially by AnimalBiome^TM^. This study aims to quantitate the colony forming units per gram (CFU/g) in lyophilized AnimalBiome^TM^ FMT products compared to fresh and lyophilized in-house FMT; and to evaluate microbial compositions across multiple FMT products. FMT products were cultured in aerobic and anaerobic environments. 16s rRNA amplicon sequencing (V4 region) was performed on FMT products and colonies taken from FMT cultures. Three lots each of AnimalBiome^TM^ DoggyBiome^TM^ (DB), DoggyBiome^TM^ from raw fed dogs (DBR), and KittyBiome^TM^ (KB) were evaluated. Freshly processed stool from screened donors enrolled in The Ohio State University Companion Animal Fecal Bank (CAFB) were used as controls. Freshly processed feces yielded significantly greater total CFU/g compared to all lyophilized products (dogs, P<0.01; cats, P<0.01). KB and feline CAFB lyophilized products exhibited comparable viability (P=0.14). Canine CAFB lyophilized FMT yielded significantly greater CFU/g than DB (P=0.17) and DBR (P=0.018). Each donor has a unique microbial profile (PERMANOVA; dogs, P=0.001; cats, P=0.03). DBR FMT products have a significantly greater abundance of *Enterobacteriaceae* compared to other canine products (P<0.01); however, no AnimalBiome^TM^ product showed detectable growth of Gram-negative microbes. These findings will provide evidence for practitioners when choosing FMT products for their patients. Further research is needed to determine the impact of FMT processing on engraftment and clinical outcome.

## 1. Introduction

The mammalian intestinal tract harbors a diverse and densely populated microbial ecosystem composed of bacteria, archaea, fungi, protozoa, and viruses. These organisms define the intestinal microbiota. Eubiosis is a diverse and balanced state of microbes that provide the host with beneficial metabolites [1]. Bacteria comprise the largest proportion of these microbial inhabitants, and thus are the major contributor to metabolic output [2].These various metabolites impact the health of the local intestinal tissue, as well as distant organs such as the brain, lungs, heart, liver, and kidneys [2]. Notable microbial metabolites include microbial-derived secondary bile acids, which aid in pathogen colonization resistance, cell signaling, and metabolic regulation [3]. Microbial-derived short-chain fatty acids are also important cell signaling molecules and contribute to intestinal motility and immune regulation [4–6]. This complex metabolic activity and importance for host health makes the gut microbiome an important immune and metabolic organ [2].

Many gastrointestinal (GI) diseases, non-GI diseases, and environmental factors are associated with intestinal dysbiosis, defined as an aberrant microbial community structure with subsequent microbial metabolic alterations. This altered microbial functional output ultimately changes the physiologic impact of the gut microbiota on the host. For example, dysbiosis in the context of diseases such as chronic kidney disease and obesity negatively effects nutrient metabolism [7–11]. In veterinary medicine, intestinal dysbiosis is associated with obesity [6], chronic enteropathy [12, 13] atopic dermatitis [14], chronic kidney disease [8], exocrine pancreatic insufficiency [15], diabetes mellitus [16], lymphoma [17], colorectal tumors [18], parasitism [19, 20], and parvoviral enteritis [21, 22]. In human medicine, intestinal dysbiosis is implicated in the pathogenesis and progression of inflammatory bowel disease [23]; and given the similarity of human and non-human mammalian microbiomes, the same is likely true in companion animals such as dogs and cats [7, 24–26].

Fecal microbiota transplant (FMT) therapy is the transfer of fecal material from a healthy donor into the GI tract of a diseased recipient to improve microbial diversity and metabolic function aimed at conferring a health benefit to the recipient. The biomass of FMT consists predominantly of bacteria, but FMT has other important constituents, including bacteriophages, fungi, protozoa, bacterial metabolites and host-derived immunoglobulins [27]. FMT has been successfully used to treat canine and feline diseases including feline ulcerative colitis [28], canine inflammatory bowel disease [29], canine chronic enteropathy [30], canine parvoviral enteritis [31], anxiety and cognitive disfunction associated with epilepsy [32], and canine atopic dermatitis [14]. Ongoing clinical trials investigating the efficacy of FMT as an adjunctive treatment for obesity and diabetes mellitus are underway [33]. The mechanism by which FMT confers a health benefit is poorly understood [33, 34], but is likely multifactorial and linked to the viability of commensal microbes, subsequent microbial engraftment, and restoration of eubiosis leading to restoration of microbial metabolic function which subsequently impacts host physiology [27, 34].

The growing clinical and public interest in FMT has prompted the development of established clinical guidelines by the Companion Animal FMT Consortium, an international consortium of veterinary gastroenterologists and FMT experts [33]. These guidelines aim to standardize fecal donor screening and FMT procedures in order to ensure consistency, safety, and efficacy for its use in companion animals[33]. The Companion Animal FMT Consortium recommends the use of FMT as an adjunctive microbial-directed therapeutic in dogs with parvoviral enteritis and acute colitis, as well as chronic enteropathy in both dogs and cats [33]. These recommendations are based on compelling evidence for the clinical efficacy of FMT in these disease conditions [29–31, 35, 36].

Growing scientific and clinical interest in FMT has also led to the development of veterinary fecal repositories, also called fecal banks. Managing an in-house fecal bank requires substantial monetary and time commitments. Fecal donors must undergo rigorous screening through a comprehensive panel of diagnostic tests to ensure the safety and quality of FMT products [33]. This thorough evaluation is essential to prevent the transmission of infectious disease or detrimental phenotypes to the FMT recipient including obesity. In addition to diagnostic testing, the Companion Animal FMT Consortium recommends the exclusion of canine and feline fecal donors if there is a history of pharmaceuticals or diets known to induce dysbiosis [33].

Following a thorough review of diet and lifestyle histories, along with comprehensive diagnostic screening, owners of fecal donors must then commit to the regular and timely delivery of fecal samples to the laboratory. Feces should be processed and used or stored within 2-6 hours of defecation for dogs, and within 24 hours for cats [33]. The Companion Animal FMT Consortium recommends that fresh feces be used whenever possible [33]; however, the narrow time window needed for delivery to the laboratory requires coordination by the practitioner and fecal donor owner, which may complicate the practical implementation of FMT in clinical practice. If using fresh feces is not feasible, the fecal material should be frozen to maintain optimal bacterial viability throughout the duration of storage [33], however this may not be possible in many veterinary and at-home settings due to biohazard concerns or resource limitations. Due to the time consuming and costly nature of managing an in-house fecal bank, many veterinarians have instead opted for commercially available FMT products for their patients to ensure consistent and convenient on-demand access to FMT products.

AnimalBiome^TM^ (Oakland, CA 94609, USA) is a privately owned company that provides veterinarians and consumers with commercially available microbiota-directed therapies for pets, including FMT. The FMT processing method used by AnimalBiome^TM^ is proprietary, however their FMT products are lyophilized and glycerol is used as the cryopreservative. Lyophilization is the process of freezing fecal material at -80°C, then freeze-drying through sublimation to achieve a dry powder that can be stored at room temperature [33]. The addition of a cryopreservative, such as glycerol, improves microbial viability and extends the shelf-life of the FMT product [33]. AnimalBiome^TM^ offers encapsulated FMT products for at-home oral administration and unencapsulated lyophilized fecal powder for reconstitution and enema administration by a licensed veterinarian. In contrast to the guidelines put forth by the Companion Animal FMT Consortium, AnimalBiome^TM^ offers FMT products from dogs exclusively fed a raw diet, in addition to FMT products from dogs and cats fed a “Standard” or conventional commercial non-raw diet. To date, the clinical efficacy of FMT products from raw fed fecal donors compared to commercially fed fecal donors remains unexplored.

The viability of microbes in FMT products plays an important role in the success of engraftment within the recipient’s GI tract [27, 34, 37, 38]. Only viable and functional microbial populations have the capacity to colonize and integrate into the gut microbiome, which is essential for restoring eubiosis and function [39]. In human medicine, the technique used to process and store FMT products impacts microbial viability [38, 40–43] and lyophilized FMT products are proven to be biologically active and a practical alternative to freshly prepared FMT [42, 44]. However, there is currently no evidence quantitating the viability of microbes in lyophilized veterinary FMT products. Given the importance of microbial viability in the proposed mechanism of action of FMT products, this study first aimed to compare and quantify the viability of microbes in commercially available AnimalBiome™ lyophilized FMT products with those from our in-house companion animal fecal bank (CAFB) lyophilized FMT products. Additionally, we compared these lyophilized FMT products to freshly processed FMT to assess the impact of lyophilization on microbial viability in canine and feline feces. Culture-dependent techniques were used to assess viability; however, due to the unique growth requirements of many intestinal microbes, only a small fraction of gut bacteria can be isolated using standard culture methods [45, 46]. Thus, the colonies successfully cultured from the FMT products tested serve as surrogates of viable microbes in these FMT products, thus, representing a subset of the broader microbial community. Further, due to the well-documented impact of a raw food diet on the canine microbiome [47–52], our secondary aim was to compare the overall microbial community structure between raw-fed and standard-fed canine fecal donors using 16S rRNA amplicon sequencing. Our hypotheses are: 1) freshly processed canine and feline FMT products will exhibit significantly greater surrogate microbial viability compared to all lyophilized FMT products regardless of source; 2) AnimalBiome^TM^ and CAFB lyophilized FMT products will have comparable surrogate microbial viability; and 3) canine fecal donors fed a raw diet will have a significantly higher abundance of *Enterobacteriaceae* compared to dogs fed a standard-diet. This foundational study is the first to provide evidence for the revivification potential, and thus functional potential, of commercially available lyophilized companion animal FMT products and to describe compositional differences in the microbiota of raw-fed and standard-fed canine fecal donors. This will inform the clinical decisions of veterinarians when choosing the most effective FMT products for their canine and feline patients.

## 2. Materials and methods

### 2.1 Fecal Donors and FMT Products

Three AnimalBiome^TM^ FMT products were tested including: DoggyBiome^TM^ Gut Restore Supplement from Standard Diet-Fed Dogs (DB), DoggyBiome^TM^ Gut Restore Supplement from Naturally Reared Dogs (DBR), and KittyBiome^TM^ Gut Restore Supplement (KB) FMT capsules. Per AnimalBiome^TM^ , DBR FMT products are derived from fecal donor dogs that are raised holistically and given a plant-based diet supplemented with raw meat [53]. AnimalBiome^TM^ collects feces from healthy donor pets that meet rigorous screening criteria to ensure high-quality and safe FMT products [53]. All animals must not have any current or prior health problems, no antibiotics administration within 6 months of donating, good fecal consistency, healthy body weight, and a good temperament [53]. Feces are regularly tested for the presence of any bacterial, viral, protist, or helminth pathogens using quantitative polymerase chain reaching (PCR) [53]. The precise FMT processing technique used by AnimalBiome^TM^ is proprietary; however, all FMT products are cryoprotected and freeze-dried[53]. Glycerol is listed as an ingredient in each FMT product; however, the concentration of this cryopreservative is unknown. Three separate lots from each FMT product, each derived from the same AnimalBiome^TM^ fecal donor, were tested for surrogate microbial viability and the microbial community structure was evaluated with 16S amplicon sequencing (Table1). Demographics of AnimalBiome^TM^ fecal donors are unknown. All FMT capsules were purchased and received directly from AnimalBiome^TM^. Upon delivery FMT capsules were stored per manufacturer instructions in their original bottle in a cool and dry environment until surrogate microbial viability testing was performed. The date of defecation, fecal collection and FMT processing for AnimalBiome^TM^ FMT products are unknown, however, all surrogate microbial viability testing was performed at least one month prior to the expiration date listed on each FMT product.

One screened CAFB canine fecal donor and a household of two screened CAFB feline fecal donors each provided naturally voided fecal samples (Table 1). Two feline fecal donors were included in this study because they reside in the same household, which ensures similar environmental and dietary exposures. Collecting fecal samples from only one cat in a multi-cat household presents logistical challenges, such as the need to isolate cats for fecal collection. Enrolling all cats in a single household facilitates consistent fecal sample collection without concern for cross-contamination of feces from unscreened donors. All CAFB fecal donors were client-owned and fed a standard commercial non-raw diet. All protocols were approved by The Ohio State University International Care and Use Committee (IACUC number 2017A00000093-R1).

**Table 1.** AnimalBiome^TM^ FMT products and lot numbers and CAFB fecal donor demographics. For each FMT product, original lot numbers are transcribed to Lot 1, 2, and 3 for simplicity. The same CAFB fecal donor provided naturally voided feces for controls in each experimental replicate. Abbreviations: FMT, fecal microbial transplant; CAFB, Ohio State University Companion Animal Fecal Bank; DB, DoggyBiome^TM^, DBR, Doggybiome^TM^ from naturally reared dogs; KB, KittyBiome^TM^; FS, female spayed.

| **AnimalBiome™**<br>**FMT Product** | **AnimalBiome™ Fecal Donors** |
| --- | --- |
| DoggyBiome™<br>Gut Restore FMT Capsules (DB) | D22112 (Lot#1)<br>D22183 (Lot #2)<br>D22184 (Lot #3) |
| DoggyBiome™<br>Naturally Reared Dog<br>FMT Capsules (DBR) | D19307 (Lot #1)<br>D19928 (Lot #2)<br>D20331 (Lot#3) |
| KittyBiome™ Gut Restore FMT<br>Capsules(KB) | C7979 (Lot #1)<br>C7984 (Lot #2)<br>C8066 (Lot #3) |
| **Companion Animal**<br>**Fecal Bank (CAFB)** | **CAFB Fecal Donor Demographics** |
| Canine Donor | 3 yo FS Miniature Dachshund |
| Feline Donors | 3.5 yo FS DSH<br>3.5 yo FS DSH |

### 2.2 Companion Animal Fecal Bank FMT Preparation

Naturally voided canine feces were refrigerated within 15 minutes of defecation and processed in the laboratory within 2 hours of defecation. Naturally voided feline feces were collected from the litterbox and processed within 12 hours of defecation. Owners were instructed to transport feces on ice in an insulated cooler to the laboratory. Feces was aerobically processed using a double centrifugation technique with 10% glycerol as previously described [54, 55]. Briefly, plant material or litter were scraped off of feces using a metal spatula. Feces were weighed and placed into a sterile Stomacher^TM^ 400 Classic Standard bag^TM^ (Seward^TM^, round bottom Stomacher^TM^ bag for Model 400, Fisher Scientific, catalog # 14-285-284). The feces were then kneaded by hand until completely homogenized. Sterile 0.9% saline was added to a ratio of 1:4 feces to sterile saline and homogenized. The saline and feces mixture were then placed into 50mL sterile conical tubes and centrifuged at 400g for 30 minutes at -4°C. Supernatant was removed to a 1:0.75 ratio of fecal sediment to supernatant. The remaining supernatant was vortex with the fecal sediment until the mixture was homogenized. The resulting fecal solution was then strained using the integrated strainer element of a sterile Stomacher^TM^ Circulator Strainer bag (Seward^TM^ Strainer bag for Stomacher ^TM^ 400, Fisher Scientific, Catalog # 14-285-22). The filtered fecal suspension was then placed into a sterile 50mL conical tube and centrifuged again at 400g for 15 minutes at 4°C. Supernatant was then removed to a 1:0.75 ratio of sediment to supernatant. The remaining supernatant was vortexed with the fecal sediment until homogenized. Glycerol was then added to a final concentration of 10% as the cryopreservative. An aliquot of the final fresh FMT slurry (CAFB Fresh) was then taken for surrogate bacterial culture. The remaining FMT slurry was frozen at -80°C for 48 hours, then lyophilized at -87.5°C and 0.005 mBar for 48 hours (FreeZone Benchtop Freeze Dryer, LabConco, Kansas City, MO, USA). Immediately upon removal from the lyophilizer, lyophilized CAFB FMT products (CAFB Lyo) were homogenized and cultured.

### 2.3 Surrogate Microbial Viability

Figure 1 demonstrates the study design and workflow to evaluate overall surrogate microbial viability and identification of viable microbes. In this study, a ‘replicate’ refers to an independent repetition of the entire culture experiment, each performed on a different day using a new aliquot from the same FMT product. Three lots each of DB and DBR FMT products were tested in two replicate culture experiments performed on separate days. Three lots of KB were tested in three replicate culture experiments performed on separate days. Freshly processed CAFB FMT product was cultured concurrently with AnimalBiome^TM^ FMT products, and lyophilized CAFB FMT were plated after removal from the lyophilizer unit. Each lot of CAFB FMT products were derived from a single defecation event. Feces were naturally voided and brought to the laboratory on the same day of each culture experiment. Plating was performed in duplicate for every culture experiment.

Aliquots of equal volume of AnimalBiome^TM^ and CAFB FMT products were diluted in sterile 1X phosphate-buffered saline (PBS) to a 1:10 stock solution, then serially diluted with 1X PBS and placed in an aerobic or anaerobic environment. All serial dilutions were performed under aerobic conditions. Three different dilutions of each FMT product were plated to ensure at least one dilution would yield a countable number of bacterial colonies on the agar plate for enumeration. Dilutions were inoculated onto MacConkey agar (Becton Dickinson Biosciences [BD] Difco^TM^) using the track plating technique as previously described [56] in order to determine the concentration of viable surrogate Gram-negative microbes, specifically *Enterobacteriaceae*. Columbia colistin naladixic acid (CNA) agar with 5% sheep blood (Remel, ThermoFisher Scientific) using the spread plate technique [57] was also performed in order to determine the concentration of viable surrogate Gram-positive microbes. Colony forming units (CFU) were enumerated after 48 hours of incubation at 37°C in both aerobic and anaerobic conditions. Anaerobic plating and incubation were performed in a vinyl anaerobic chamber at 0-5 ppm oxygen and 2.0 to 2.5% hydrogen concentration (Coy Laboratory Products, Grass Lake MI). Agar plates were placed in the anaerobic chamber 12 hours prior to inoculation.

To determine the identity of the surrogate viable microbes, visible bacteria colonies were collected from agar plates for 16S rRNA gene amplicon sequencing and analysis (Figure 1). Briefly, after 48 hours of incubation, one hundred colonies were individually scraped from aerobic agar plates using a sterile inoculating loop and placed in a sterile 2mL epitube filled with 500 uL sterile 1X phosphate buffered saline (PBS). Using the same technique, an additional 100 colonies were collected from anaerobic plates and placed in a separate sterile epitube filled with 500 uL sterile 1X PBS. Every unique colony morphology identified on all agar plates were collected using this method. Following colony collection, all epitubes were immediately frozen at -80°C for storage. Epitubes were thawed on ice, homogenized, and combined prior to submission for DNA extraction and amplicon sequencing. Colony collection was performed for all experiments except one feline culture experiment.

**Figure 1.**
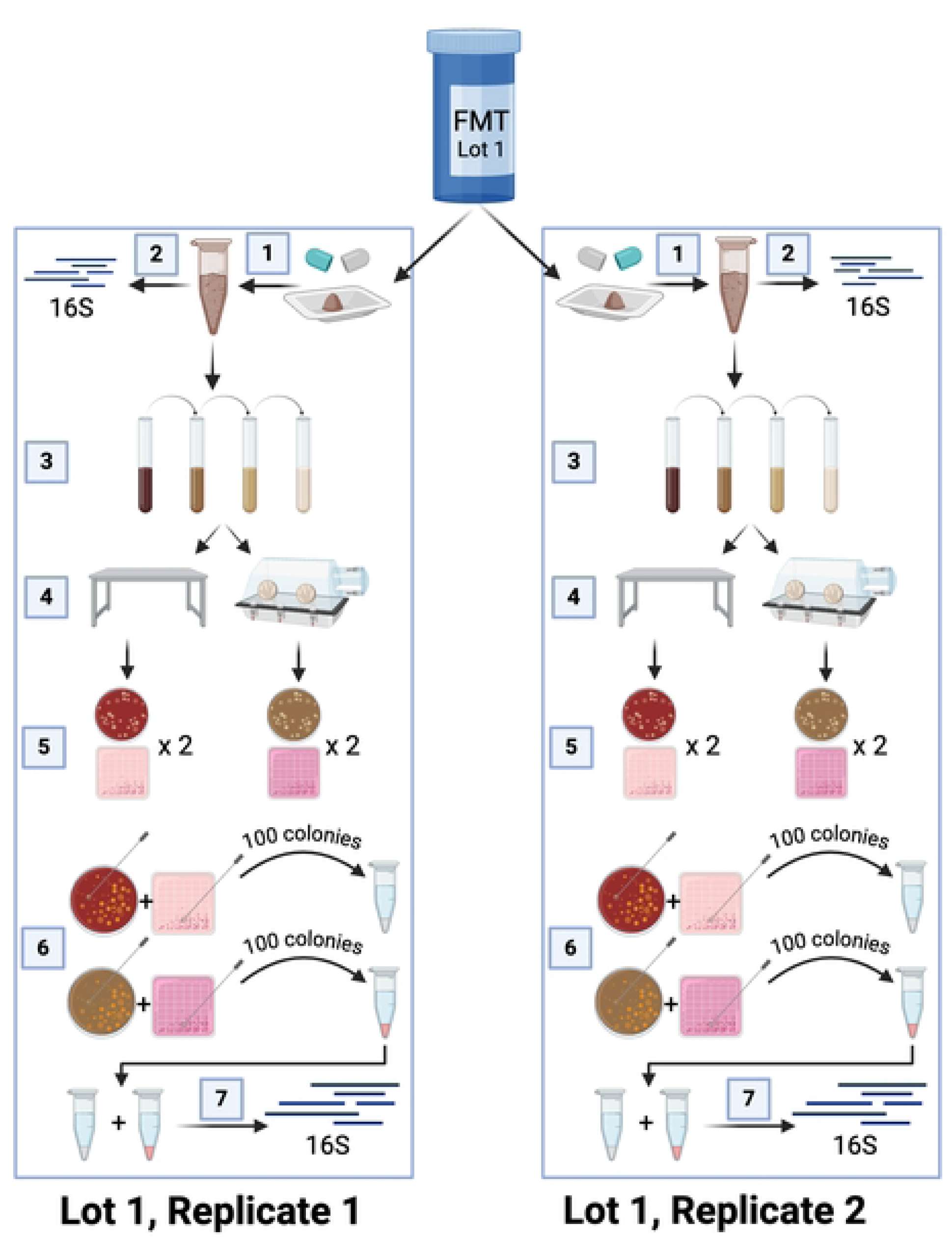
Schematic of the study design and workflow for microbial viability testing and 16S rRNA gene amplicon sequencing of companion animal FMT products and collected microbial colonies. 1) AnimalBiome^TM^ FMT capsules were opened under a sterile hood and their contents were weighed. 2) One aliquot of the FMT material was stored at -80℃ until DNA extraction and sequencing. 3) One aliquot was serially diluted under aerobic conditions. 4) Serial dilutions were cultured in both aerobic and anaerobic conditions 5) Dilutions were plated on two selective agars: CNA Blood Agar for Gram-positive microbes and MacConkey agar for Gram-negative microbes. Agar plates were incubated at 37℃ for 48 hours. 6) 100 colonies were selected from the aerobic and anaerobic plates and placed into sterile epitubes containing 500 uL 1XPBS 7) The combined samples were submitted for DNA extraction and subsequent 16S rRNA gene amplicon sequencing. Figure generated using BioRender.com.

### 2.4 DNA Extraction and Amplicon Sequencing

Every lot from each FMT product underwent DNA extraction and amplicon sequencing at the University of Michigan Microbiome Core. DNA was extracted from FMT products and surrogate viable microbes using a MagAttract PowerMicrobiome kit (Qiagen, Germantown, MD, USA) following manufacturer instructions. Sequencing of the V4 region of the 16S rRNA gene amplicon was performed using a dual indexing approach as previously described[58, 59]. Briefly, genes were amplified using universal primers 515f and 806r[60]. DNA products were initially incubated at 97°C for 120 seconds; followed by 30 PCR amplification cycles. Denaturation occurred at 95°C for 20 seconds, annealing at 55°C for 15 seconds, then 72°C for 900 seconds. PCR products were maintained at 4°C until product visualization with SYBR Safe DNA Gel Stain (ThermoFisher Scientific). Library normalization was performed using a SequalPrep Normalization Plate Kit (Life technologies, Carlsbad, CA, USA). The final concentration of pooled samples was performed using a Kapa Biosystems Library Quantification kit for Illumina platforms (Kapa Biosystems, Wilmington, MA, USA). Samples were normalized to the lowest sample concentration.

Amplicon sequencing was performed using an Illumina MiSeq platform with a MiSeq reagent kit (Illumina, San Francisco, CA, USA) using V2 chemistry with 500 cycles as previously described [59]. FASTQ files were generated for each paired-end read. Master mix and sterile extraction reagents served as negative controls throughout DNA processing, amplification, and sequencing procedures.

### 2.5 Microbiota Analysis

Amplicon (V4 16S rRNA gene) sequences were imported into R Studio (Version 2023.09.1+494, Vienna, Austria) and paired-end reads were assembled into contigs, trimmed, filtered, and converted into amplicon sequence variants (ASV) using the DADA2 pipeline as previously described [61]. Chimera sequences and amplicon sequence variants (ASVs) <250 base pairs or >256 base pairs in length were removed. Fastq files from KittyBiome^TM^ Lot 8066 (Lot 3) and DoggyBiome^TM^ Lot D22184 (Lot 3) were excluded prior to statistical analysis due to low read depth (<5,000 sequences per fastq file). The low read depth was unexpected for fecal samples due to the high bacterial biomass present in feces. This may be due to problems with sample acquisition, sample storage, and/or DNA extraction, rather than a true reflection of microbial abundance.

Taxonomic classification of ASVs and taxonomy table generation was performed using the phyloseq package (Version 1.40.0) in R Studio with the Silva 16S rRNA Sequence Database (Version 138.1) [62, 63]. ASVs that comprised less than 1% of the total sequences in each sample were removed from composition bar plots and listed as “other” for data visualization. Alpha and beta diversity analysis were performed using the phyloseq and vegan (Version 2.6-2) packages in R studio (Version 2023.09.1+494, Vienna, Austria).

### 2.6 Statistical Analysis

Alpha diversity metrics was assessed using total observed ASVs and Shannon Index. For alpha diversity, statistical significance was evaluated with the One-Way Analysis of Variance (ANOVA) with corrections for multiple comparisons applied using the two-stage linear step-up procedure of Benjamini, Krieger, and Yekutieli. Due to the small sample size for feline dataset, only descriptive statistics were reported.

Beta diversity of FMT products was assessed using the phyloseq package in R Studio, employing the Bray-Curtis dissimilarity metric. Statistical significance of beta diversity was determined through Permutational Analysis of Variance (PERMANOVA). Visualization of beta diversity was achieved through principal coordinate analysis (PCoA). To assess the variability of the microbial communities within each donor group, the betadisper() function from the vegan package was used to calculate the Bray-Curtis dissimilarity distances between each individual samples and the centroid of its respective donor group. Using this metric, larger distances indicate more dissimilar microbial community structures. Statistical significance of Bray-Curtis distances from the centroid were tested using the Kruskal-Wallis test, followed by post hoc comparisons using the Benjamini, Krieger, and Yekuteli procedure. Statistical comparisons of alpha diversity and dispersion distances were performed using GraphPad Prism (Version 9.4.0, GraphPad Software, LLC, La Jolla, CA, USA). For the amplicon sequencing data from the collected viable aerobic and anaerobic colonies, only the “presence/absence” of surrogate viable microbial families was assessed.

A consensus of differentially abundant microbial families across standard-fed and raw-fed canine FMT products was obtained using three differential abundance tests, Linear discriminant analysis Effect Size (LEfSe), Microbiome Multivariable Associations with Linear Models (MaAsLin2), and Analysis of Compositions of Microbiomes with Bias Correction (ANCOM-BC). LEfSe was conducted by rarefying the data to 22,608 sequences, corresponding to the lowest read depth across all canine FMT products. The data was then normalized using trimmed mean of M-values (TMM). All data were normalized using total sum scaling (TSS) prior to analysis with MaAsLin2. LEfSe and MaAsLin2 were performed using MicrobiomeAnalyst [64]. ANCOM-BC was performed using the ANCOMBC package (Version 2.6.0) in R Studio. The model included the experimental condition “diet” as a fixed effect. To account for multiple testing, p-values were adjusted using the Holm method, and a significance level of 0.05 was applied.

Differential abundance testing was also used to compare the microbial compositions of feline CAFB and KB FMT products. LEfSe and MaAsLin2 were performed using MicrobiomeAnalyst [64]. LEfSe was conducted by rarefying the data to 23,465 sequences. The data was then normalized using TMM. For MaAsLin2, data were normalized using TSS. ANCOM-BC was not performed on feline data due to small sample size.

For each FMT product type, absolute bacterial counts (CFU/g) were calculated by totaling CFU/g from each dilution/plate. Normality testing was performed using the Shapiro-Wilk test and visualized with QQ plots. Significance testing was performed using the Ordinary One-Way ANOVA for parametric data or Kruskal-Wallis with two-stage linear step-up procedure of Benjamini, Krieger, and Yekutieli, for non-parametric data, as appropriate. For the two-stage linear step-up procedure of Benjamini, Krieger, and Yekutieli, a significant discovery was defined as a *q* value less than the corresponding *Q* value. For all other statistical testing, significance was defined as *P*<0.05.

### 2.7 Data Availability

All amplicon sequences included herein are publicly available via the National Center for Biotechnology Information Sequence Read Archive (NCBI SRA) via the following BioProject identification number: PRJNA1227492.

## 3. Results

### 3.1. Microbiota Analysis of Canine FMT Products

To evaluate the microbial community members of the FMT products, amplicon sequencing of the FMT products was performed. Within CAFB FMT products, there was no significant difference in alpha diversity metrics (Observed ASVs and Shannon Index) between fresh, denoted as “CAFB Fresh” and lyophilized FMT, denoted as “CAFB Lyo” (Observed ASVs, *P=*0.22; Shannon Index, *P=*0.27; Figure 2). DB FMT products had a significantly greater alpha diversity indices (observed ASVs and Shannon Index) compared to CAFB Lyo, DBR, and CAFB Fresh FMT products (Ordinary one-way ANOVA with two-stage linear step-up procedure of Benjamini, Krieger, and Yekutiel: Observed ASVs, *P*<0.01 to *P*=0.02 for all comparisons; Shannon Index, *P*<0.01 to *P*=0.02 for all comparisons) (Figure 2A and 2B). DB products exhibited the highest median observed ASVs (167; range: 136-194) compared to CAFB Fresh (117; range: 114-137), CAFB Lyo (100; range: 95-104), and DBR products (111; range: 95-174). The median Shannon Index was also highest in DB products (3.70; range: 3.55-3.80) compared to CAFB Fresh (3.20; range: 2.90-3.28), CAFB Lyo (3.04; range: 2.84-3.12), and DBR products (3.07; range: 2.62-3.66).

**Figure 2.**
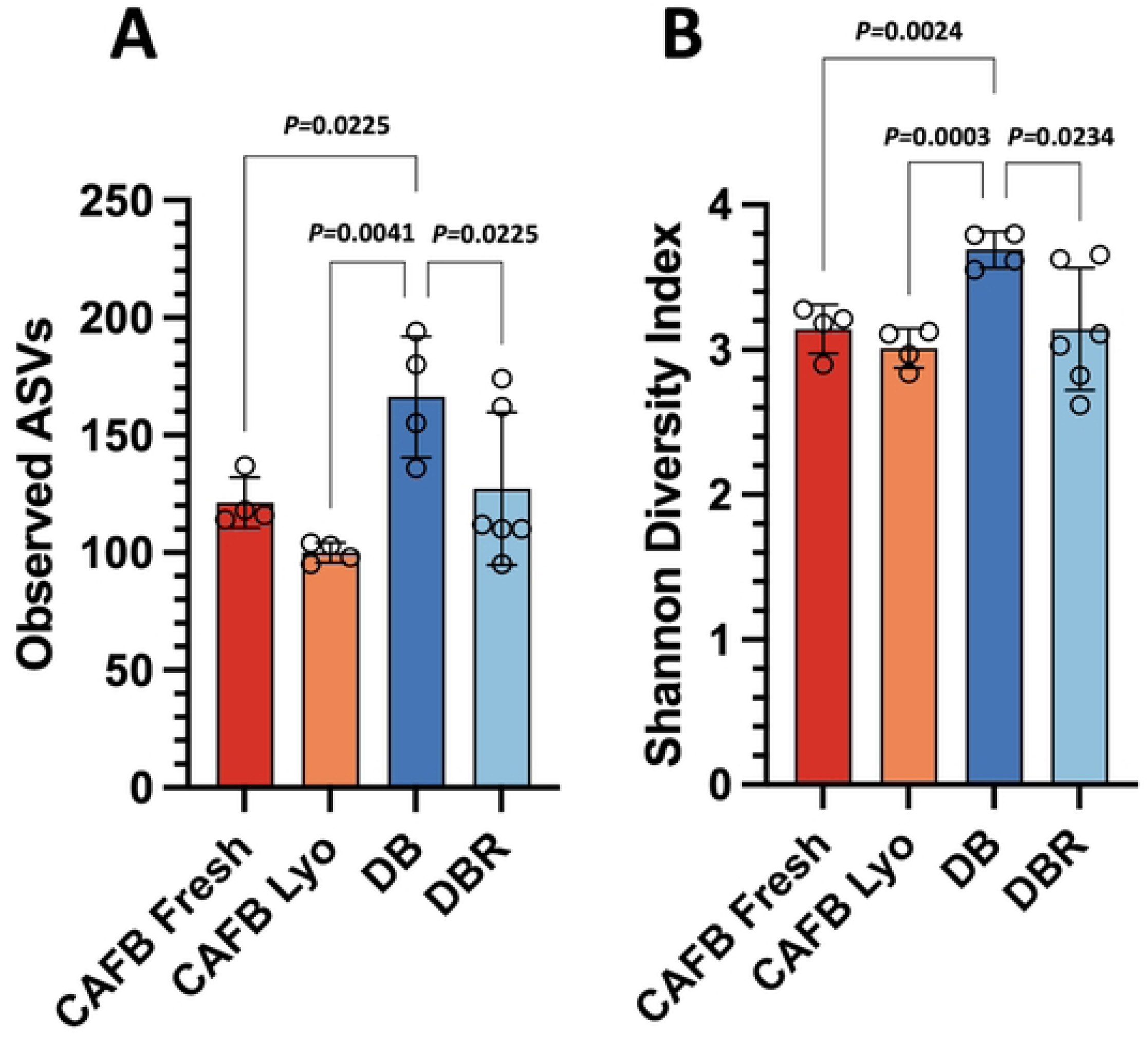
Alpha Diversity metrics for canine FMT products. DB FMT products exhibit significantly greater observed ASVs (A) and Shannon index (B) compared to the CAFB Fresh (Observed ASVs, *P*=0.02; Shannon index, *P*<0.01), CAFB Lyo (Observed ASVs, *P<*0.01; Shannon Index, *P<*0.01), and DBR FMT products (P=0.02). Significance testing was performed using the Ordinary one-way ANOVA Test with two-stage linear step-up procedure of Benjamini, Krieger, and Yekutieli.

As expected, beta diversity analysis for canine FMT products revealed that the microbial community structure varies significantly between individual fecal donors regardless of source (Bray-Curtis Dissimilarity Metric, PERMANOVA, *P*<.01 for all canine comparisons). CAFB FMT products, derived from a single FMT donor but separate defecations, form a discrete cluster and there is no significant difference in the microbial community structure between fresh and lyophilized CAFB FMT products (Bray-Curtis; PERMANOVA, P=0.61; Figure 3A). To quantitate the variability between each lot of FMT products for each individual donor, the distance of each sample from the centroid of its respective fecal donor group was calculated using the Bray-Curtis dissimilarity metric. In this metric, longer distances indicate more dissimilar microbial community structures. The median distance from the centroid for DBR FMT samples was 0.33 (range: 0.20-0.59), higher than that of CAFB Fresh (0.20; range: 0.13-0.28), CAFB Lyo (0.24; range: 0.20-0.33) and DB (0.14; range: 0.12-0.16) FMT products. However, statistically significant differences in distance from the centroid were only identified between DB and DBR (Kruskal-Wallis; *P=*0.02).

**Figure 3.**
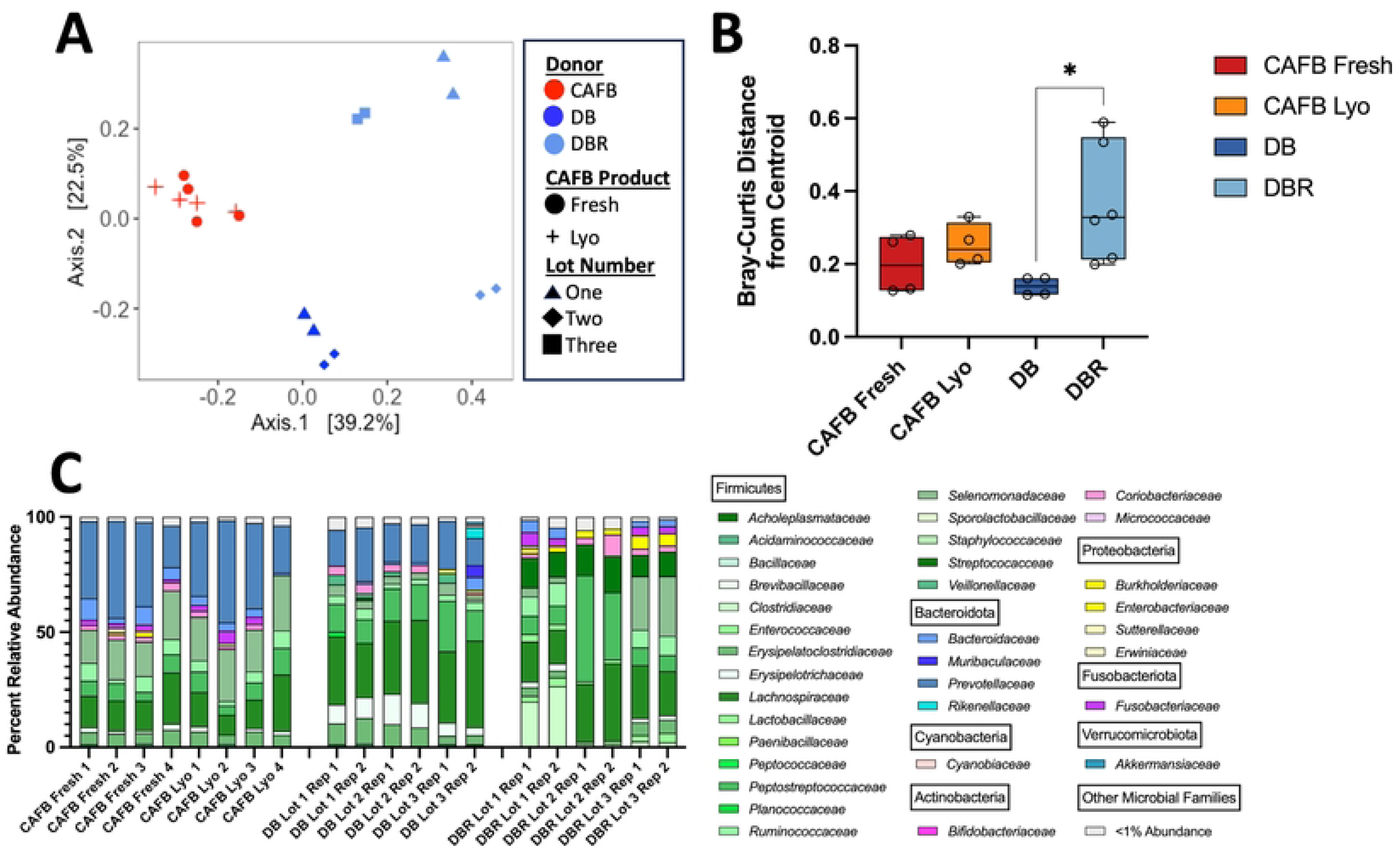
Beta diversity metrics in canine FMT products. Lot 3 of DB products were removed prior to analysis due to low read depth. A) Principal coordinate analysis of canine FMT products revealed that the microbial community structure clusters by donor of origin (Bray-Curtis Dissimilarity, PERMANOVA P<0.01 for all donor comparisons). B) The Bray-Curtis median distance from the centroid of DBR products is consistently higher than CAFB Fresh and CAFB Lyo products, however statistically significant differences were only observed between DBR and DB (P=0.02). Significance testing was performed using the Ordinary one-way ANOVA Test with two-stage linear step-up procedure of Benjamini, Krieger, and Yekutieli. A q value less than the corresponding Q value indicates discovery and are annotated with a single black asterisk. No asterisk indicates no discovery. C) The percent relative abundance of microbial phyla and families in canine FMT products. Abbreviations: CAFB, Companion Animal Fecal Bank; Lyo, lyophilized; DB, DoggyBiome^TM^; DBR, DoggyBiome^TM^ Raw-Fed.

Differential abundance testing was performed to compare canine fecal donors fed a commercial diet (CAFB and DB) to those fed a raw diet (DBR). Three differential abundance methods were performed and a consensus between the statistical methods was obtained in order to ensure biological relevance of differentially abundant microbes detected. MaAsLin2, ANCOM-BC, and LEfSE identified 8, 31, and 7 differentially abundant families, respectively. Across these three testing methods, a consensus of six differentially abundant microbial families was detected. Families grouped under the phylum Firmicutes represented four of six differentially abundant families. Standard fed dogs had higher abundance of *Prevotellaceae* compared to DBR. Five microbial families were enriched in DBR including: *Clostridiaceae, Enterococcaceae, Lactobacillaceae, Streptococcaceae, and Enterobacteriaceae* (Figure 4).

**Figure 4.**
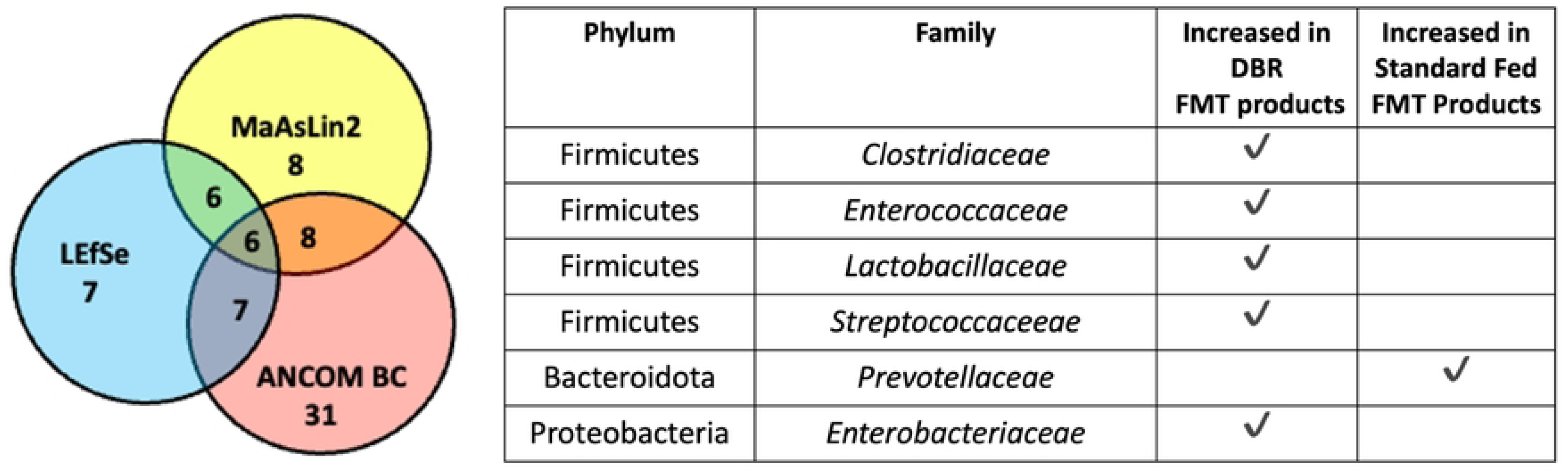
Differentially abundant microbial families in canine FMT derived from commercially fed (CAFB and DB) and raw fed dogs (DBR only). A consensus of seven microbial families was obtained using LEfSE, MaAsLin2, and ANCOM BC.

### 3.2 Microbiota Analysis of Feline FMT Products

Descriptive statistics are provided for feline alpha diversity metrics (Observed ASVs and Shannon Index) as small sample size precluded significance testing. Feline CAFB fresh FMT products exhibited higher median observed ASVs (192.5, range: 179-206) compared to CAFB Lyo (166.0; range: 157-175) and KB products (162.8; range: 147-179) (Figure 5A). The median Shannon Index was higher in CAFB Fresh (3.70; range: 3.48-3.90) and KB (3.70; range: 3.53-3.81) FMT products compared to CAFB Lyo (3.22; range: 2.92-3.51) (Figure 5B).

**Figure 5.**
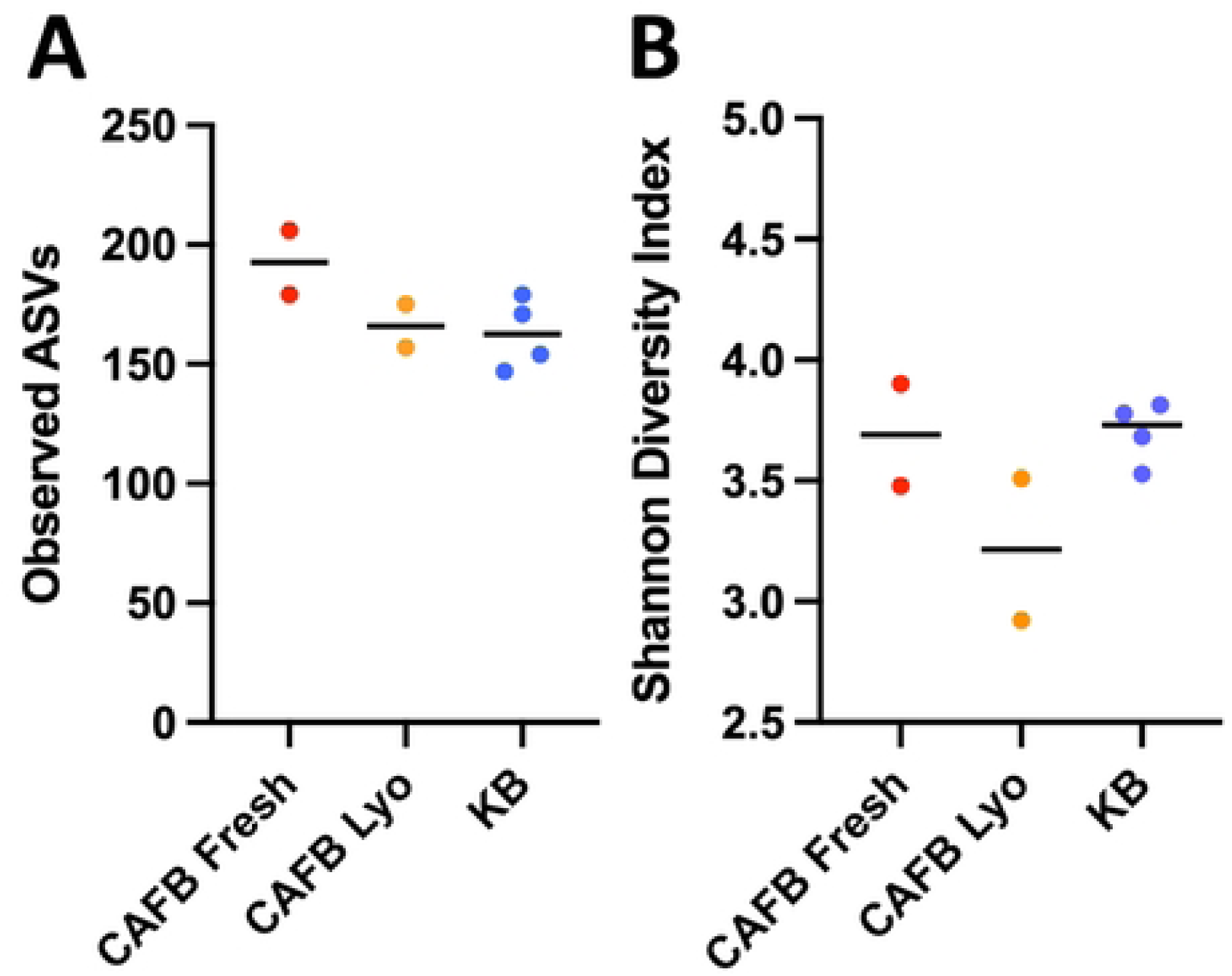
Alpha diversity metrics in feline FMT products. A) Fresh FMT products exhibited the greatest observed ASVs (206, 179) compared to CAFB Lyo (157, 175), and KB (147, 179, 171, 154) products.

Beta diversity analysis revealed that the microbial community structure varies significantly between individual donors (Bray-Curtis: PERMANOVA, P=0.03) (Figure 6A). The median Bray-Curtis distance from the centroid of CAFB Lyo and KB products was similar at 0.30 (range: 0.13-0.46) and 0.25 (range: 0.06-0.39), respectively. CAFB Fresh products had a lower median distance (0.15; range: 0.10-0.19) compared to both lyophilized products (Figure 6B). Statistical comparisons were not performed due to small sample size.

**Figure 6.**
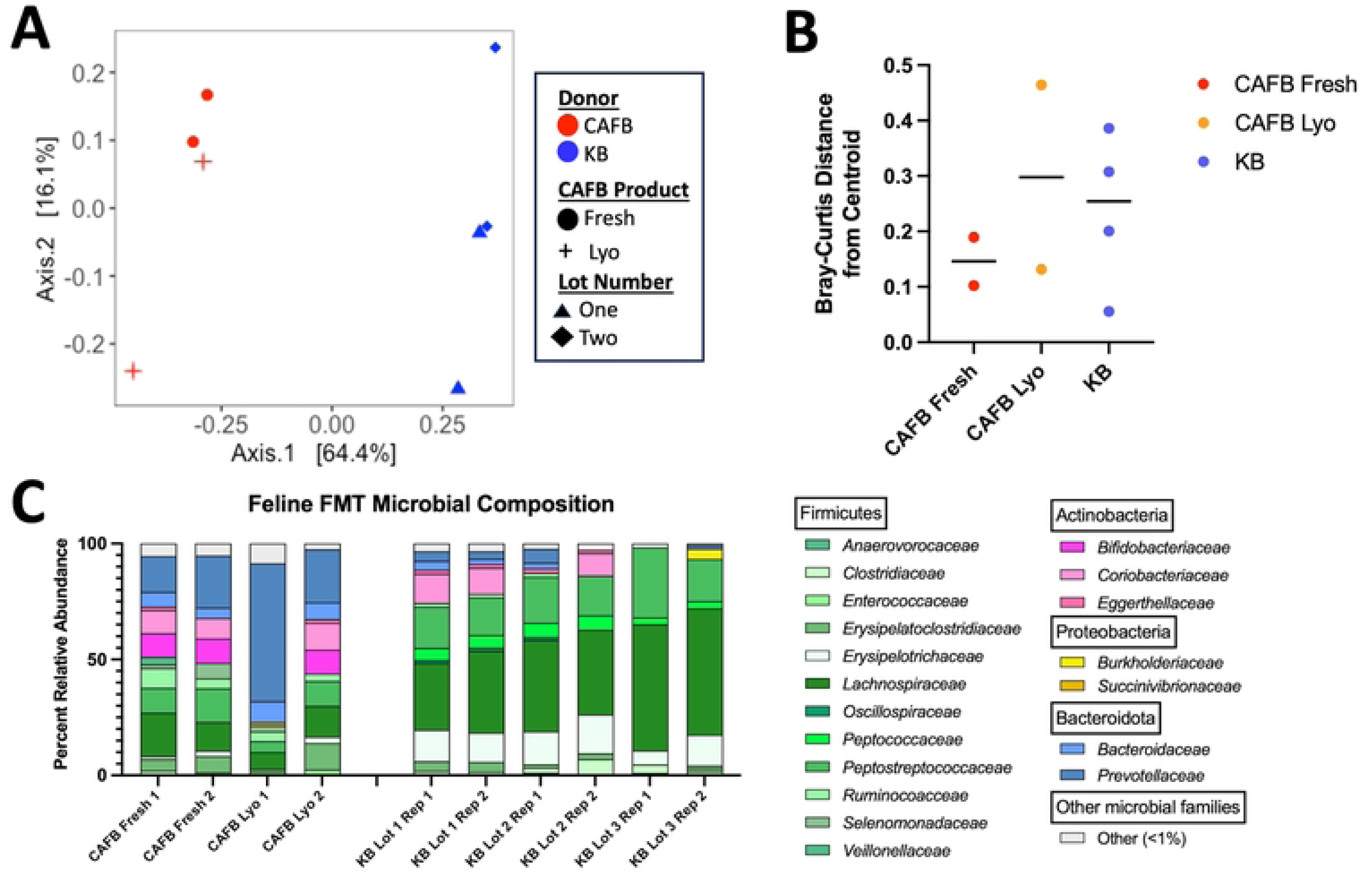
Beta diversity metrics in feline FMT products. A) Principal coordinate analysis of feline FMT products revealed that the microbial community structure varies between individual donor (PERMANOVA, P=0.03). B) The Bray-Curtis distance from the centroid of each donor group was calculated. The median distance of CAFB Lyo and KB products were similar at 0.30 and 0.25, respectively. The median distance of CAFB Fresh to its respective centroid was 0.15. Small sample size precluded significance testing. C) Relative abundance barplots of microbial families within feline FMT products. Abbreviations: CAFB, Companion Animal Fecal Bank; Lyo, lyophilized; KB, KittyBiome^TM^.

Figure 6C shows the relative abundances of microbial phyla and families in feline FMT products. Four phyla were identified across all feline FMT products. In CAFB fresh feline FMT products, Firmicutes was the most abundant phylum, ranging from 50.75-53.43%. Bacteroidota was the second most abundant phylum, ranging from 22.20-26.87%. Actinobacteria and Proteobacteria comprised 20.21-21.64% and 1.22-1.25% relative abundance, respectively. In one CAFB Lyo replicate (Feline CAFB Lyo Lot 1), Bacteroidota was the most abundant phylum (68.73%) followed by Firmicutes (25.60%). In the second CAFB Lyo replicate, Firmicutes was the most abundant phylum ranging at 46.25%, followed by Bacteroidota at 29.81%. Actinobacteriota comprised 2.08-23.21% of the microbial community. Proteobacteria comprised 2.51% of CAFB Lyo Lot 1 and was not observed in CAFB Lyo Lot 2.

Firmicutes was the most abundant phylum in all samples of KB FMT, ranging from 76.74-98.92% Bacteroidota was the second most abundant phylum in two samples, ranging from 1.59-8.52%. Actinobacteria was the second most abundant phylum in three samples, ranging from 2.34-14.80%. Proteobacteria was only observed in one sample (KB Lot#3) at 4.37% relative abundance. Nineteen microbial families were identified across all feline FMT products (Figure 6C). *Lachnospiraceae* and *Peptostreptococcaceae* were consistently present in all feline FMT products, while *Selenomonadaceae, Veillonellaceae, Bifidobacteriaceae*, and *Succinivibrionaceae* were unique to the feline CAFB products. Conversely, *Peptococcaceae*, *Oscillospiraceae*, and *Anaerovorocaceae* were exclusive to the KB FMT product.

To evaluate statistically significant differential abundance, two differential abundance methods were used. LEfSe and MaAsLin2 identified four and seven differentially abundant microbial families, respectively. Due to limited sample size, ANCOM-BC analysis was not performed in the feline FMT products. A consensus identified four differentially abundant microbial families. *Bifidobacteriaceae, Succinivibrionaceae,* and *Selenomonadaceae* were increased in feline CAFB FMT, while *Peptococcaceae* was more abundant in the KB FMT product.

### 3.3 Surrogate microbial viability

#### Surrogate microbial viability in canine FMT products

All lots of freshly processed canine FMT exhibited robust growth of aerobic and anaerobic Gram-negative and Gram-positive microbes (Figure 7A). Aerobic and anaerobic Gram-negative CFUs were detected in only one of the four lots of the CAFB Lyo FMT product, but were absent in all lots and replicates of DB and DBR FMT products (Figure 7A).

**Figure 7.**
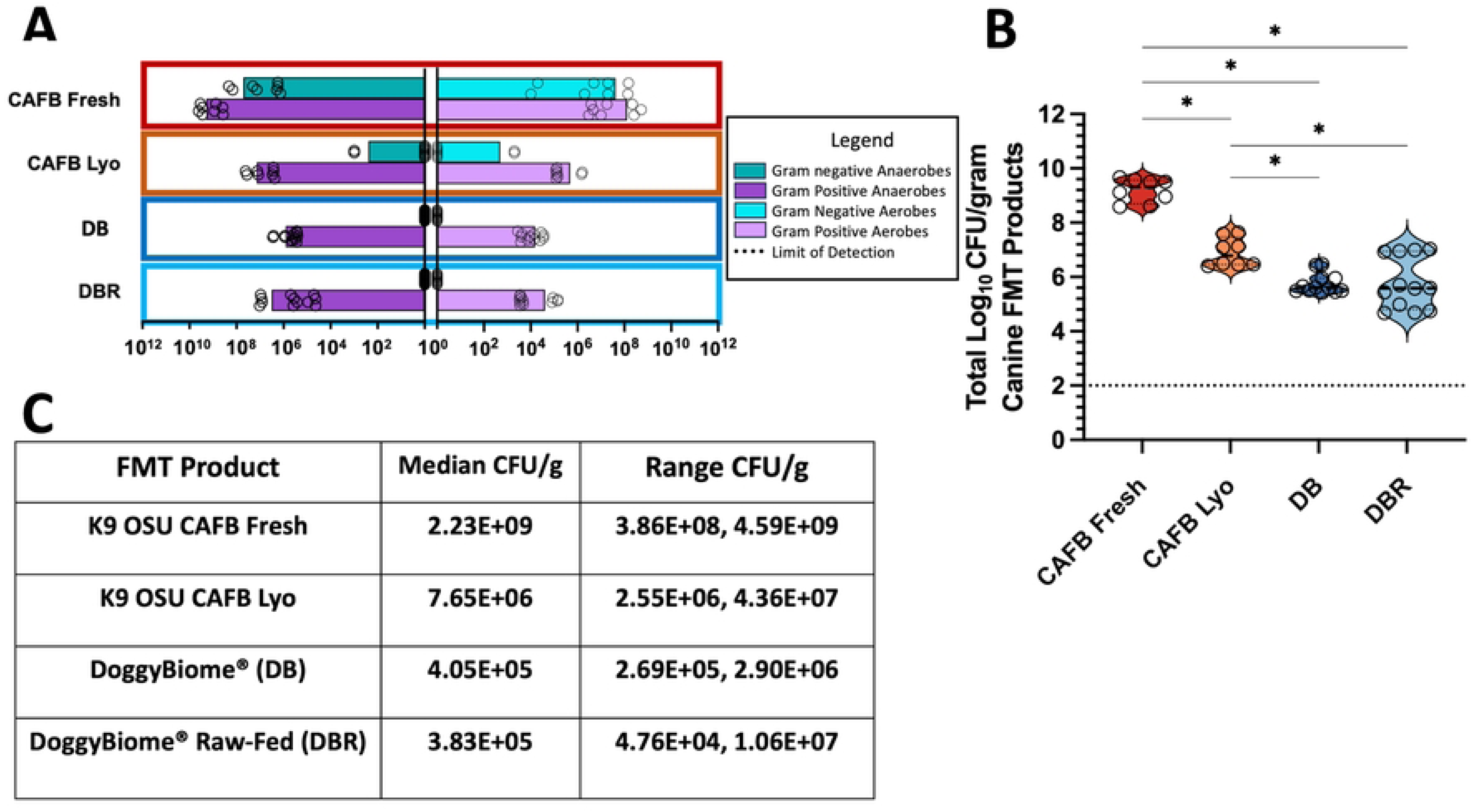
The surrogate microbial viability of in-house and commercially available canine FMT products. A) Lyophilization has a significant impact on overall microbial viability, particularly Gram-negative microbes. No viable Gram-negative microbes were observed in DB or DBR FMT products. B) Freshly processed canine FMT exhibit significantly greater microbial viability compared to CAFB Lyo and AnimalBiome^TM^ products. CAFB Lyo products exhibit statistically significantly greater CFU/g compared to AnimalBiome^TM^ products with no difference between DB and DBR. All FMT products exhibited variation in total CFU/g between individual lots. C) Median and range of total CFU/g in all canine FMT products. Significance testing was performed using Kruskal-Wallis Test with two-stage linear step-up procedure of Benjamini, Krieger, and Yekutieli. A q value less than the corresponding Q value indicates discovery and are annotated with a single black asterisk. No asterisk indicates no discovery. Abbreviations: CAFB, Companion Animal Fecal Bank; Lyo, lyophilized; DB, DoggyBiome^TM^; DBR, DoggyBiome^TM^ Raw-Fed.

As expected, freshly processed canine FMT displayed significantly higher overall surrogate microbial viability compared to all lyophilized FMT products including CAFB Lyo (Kruskal-Wallis; *P*=0.03), DB (Kruskal-Wallis; *P*<0.01), and DBR (Kruskal-Wallis; *P*<0.01) FMT products (Figure 7B). The median total CFU/g in canine CAFB Fresh FMT products was 2.23×10^9^. Canine CAFB Lyo FMT products exhibited statistically significantly greater CFU/g (median=7.65×10^6^) compared to DB (Kruskal-Wallis; *P*=0.02) and DBR (Kruskal-Wallis; *P*=0.02) products (Figure 7B). The median CFU/g of DB and DBR products were 4.05×10^5^ and 3.83×10^5^, respectively. All canine FMT products showed some variability in microbial viability between individual lots; with DBR exhibiting the largest range of CFU/g (4.76×10^4^ – 1.06×10^7^) (Figure 7C).

To determine the identity of surrogate viable microbes, 100 representative colonies were removed from aerobic agar plates and another 100 colonies from the anaerobic agar plates. These were then sequenced using the V4 region of the 16S rRNA gene. The identities of surrogate viable microbes in canine FMT products are shown in Figure 8. A total of 18 microbial families and 26 genera were identified from agar plates across all FMT products. Viable *Enterococcus* spp. was present in all lots of CAFB fresh and CAFB Lyo products, and two out of three lots of DB and DBR. *Paeniclostridium* spp was isolated in all lyophilized FMT products including CAFB Lyo, DB, and DBR, but was not present in any canine CAFB fresh FMT product. *Clostridium sensu stricto* was a viable microbe sequenced from in all DB and DBR lots but was not isolated from any CAFB FMT products. *Bacillus* spp was no isolated from any CAFB FMT products, but was present in all lots of DBR and two lots of DB FMT products. *Escherichia-Shigella* were present in all lots of CAFB fresh and one out of the four lots of CAFB Lyo; and was not identified in any DB or DBR FMT products. Microbes within Actinobacteriota, including *Bifidobacterium* spp and *Collinsella* spp were isolated from CAFB Fresh and CAFB Lyo FMT products, but were not isolated from DB or DBR FMT products.

**Figure 8.**
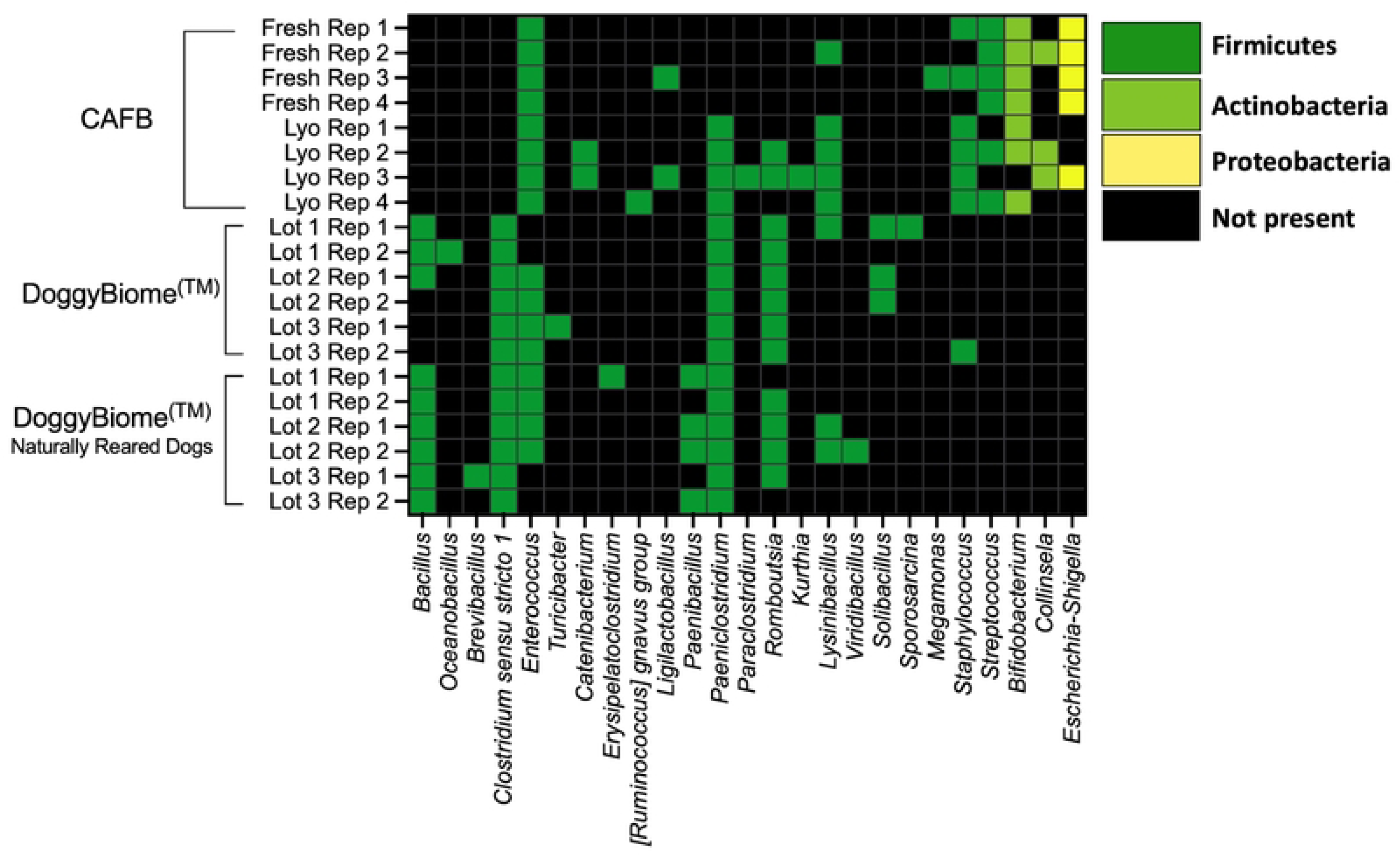
The identity of viable surrogate bacteria grown using selective agar in aerobic and anaerobic environments from canine FMT products. The diversity of viable surrogate microbes varies across donor. Dark green, light green, and yellow cells indicate the presence of their respective phyla as colonies on agar plates. Black boxes indicate no colonies were isolated from that FMT product. Abbreviations: CAFB, Ohio State University Companion Animal Fecal Bank; Lyo, lyophilized.

#### Surrogate microbial viability in feline FMT products

Overall, surrogate microbial viability in feline FMT products followed a similar pattern to canine FMT products. All lots of CAFB fresh FMT products exhibited growth of aerobic and anaerobic Gram-positive and Gram-negative microbes. Growth of Gram-negative colonies was observed in all lots of CAFB fresh FMT and only lot of CAFB Lyo. Gram-negative growth was not observed on agar plates from any KB FMT products, regardless of lot (Figure 9A).

**Figure 9.**
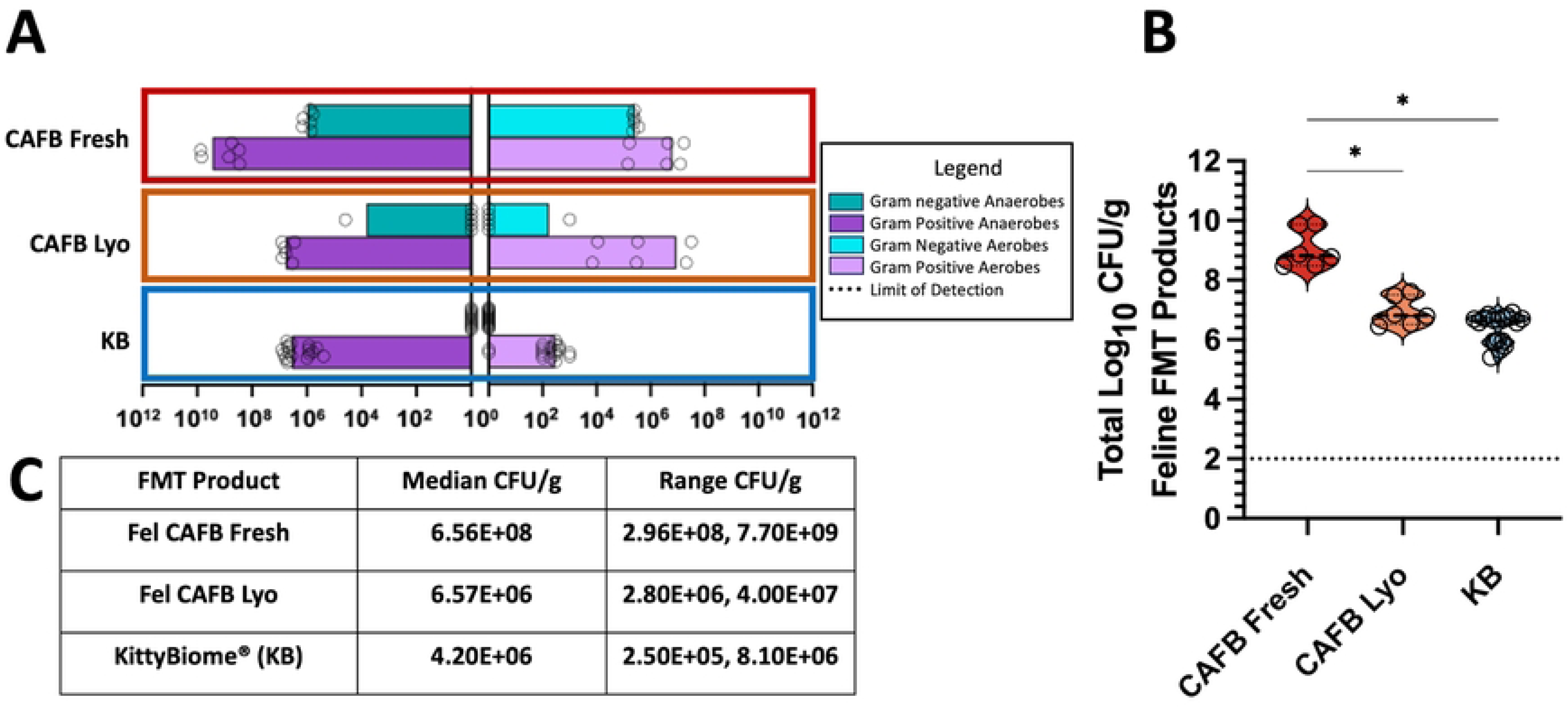
The surrogate microbial viability of CAFB and commercially available feline FMT products. A) Lyophilization has a significant impact on overall microbial viability, particularly Gram-negative microbes. No viable Gram-negative microbes were observed in KB products. B) Freshly processed feline FMT exhibits significantly greater microbial viability compared to CAFB Lyo (P=0.04) and KB products (P=<0.01). OSU CAFB feline lyophilized products exhibit comparable surrogate microbial viability compared to KB (P=0.14). C) Median and range of total CFU/g in all feline FMT products. Significance testing was performed using Kruskal-Wallis Test with two-stage linear step-up procedure of Benjamini, Krieger, and Yekutieli.

Freshly processed feline FMT exhibited significantly higher median total CFU/g compared to CAFB Lyo (Kruskal-Wallis; *P*=0.04) and KB products (Kruskal-Wallis; *P*<0.01). For the lyophilized FMT products, no significant difference was found in the overall viability between CAFB Lyo and KB (Kruskal-Wallis; *P*=0.14) (Figure 9B). The median total CFU/g of CAFB Fresh products was 6.56×10^8^ CFU/g (Figure 9C). The median total CFU/g of CAFB Lyo and KB FMT products were 6.56×10^6^ and 4.20×10^6^ CFU/g, respectively (Figure 9C).

The identities of surrogate viable microbes in feline FMT products are shown in Figure 10. Four bacterial phyla, fourteen bacterial families, and 17 genera were isolated from agar plates. Firmicutes was the most abundant phyla and was isolated from all feline FMT products, regardless of processing method. *Enterococcus* was present in all feline CAFB FMT products, and KB Lot 2. *Bifidobacterium* spp. and *Plesiomonus* spp. were present only in CAFB fresh FMT products and not in the CAFB Lyo or KB FMT products. *Clostridium sensu stricto* 1 was viable in all KB replicates and one lot of CAFB Lyo; but was not isolated in from either lot of CAFB Fresh FMT products. Similar to canine FMT products, Actinobacteria and Proteobacteria were only observed in feline CAFB Fresh and CAFB Lyo FMT products and were not isolated from any lot of KB products. *Bacillus* spp*,, Turicibacter* spp., and *Paenibacillus* spp, were isolated in at least one out of two replicate culture experiments for each lot of KB, but were not isolated from any lots of feline CAFB Fresh or CAFB Lyo FMT products.

**Figure 10.**
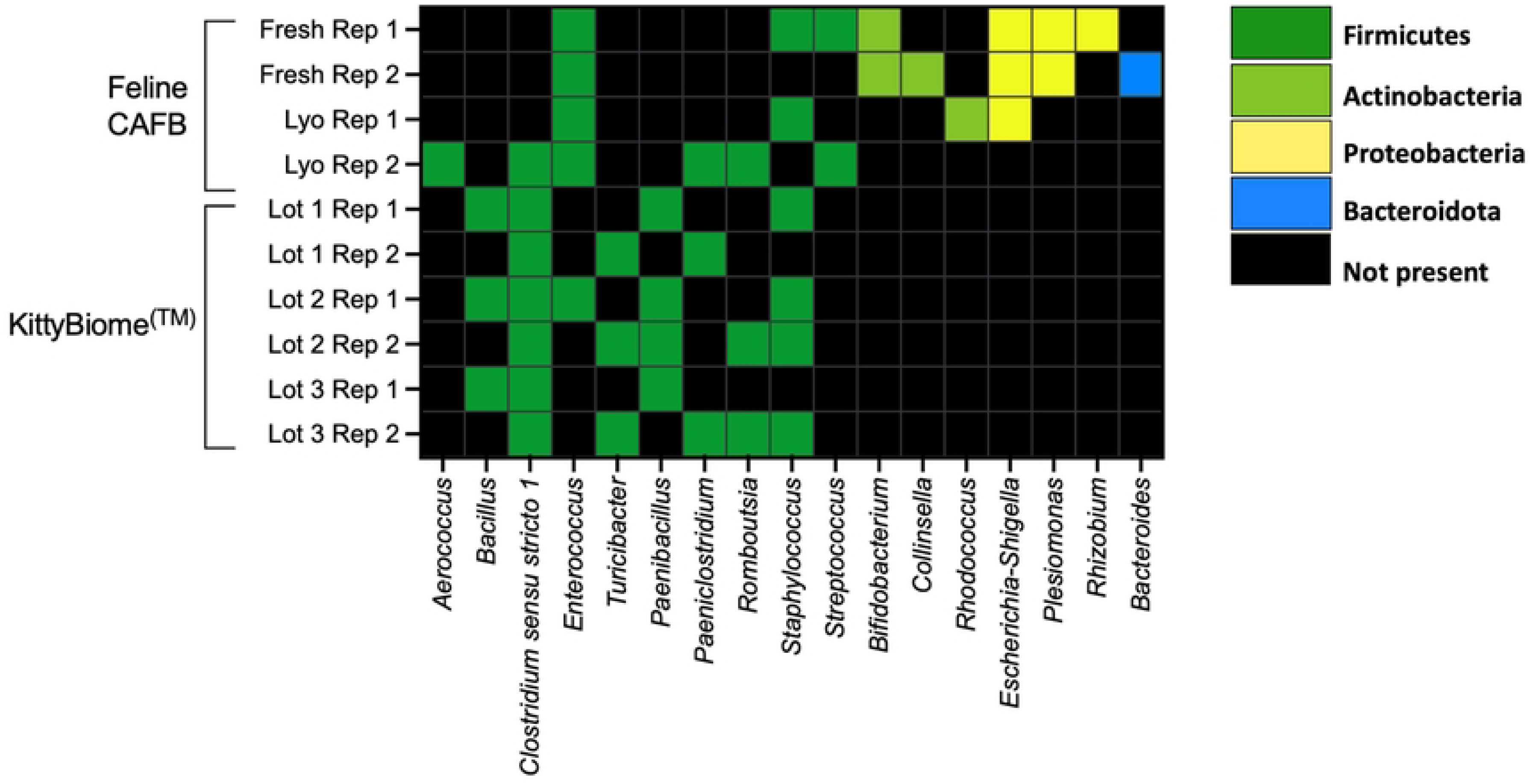
The identity of surrogate viable microbes in feline FMT products. Microbial colonies were collected from CNA and MacConkey agar following 48 hours of incubation in aerobic and anaerobic environments. Freshly processed feline OSU CAFB products exhibit greater diversity of viable microbial phyla compared to KB products. Dark green, light green, yellow, and blue cells indicate the presence of their respective phyla as colonies on agar plates. Black boxes indicate no colonies were isolated from that FMT product. Abbreviations: OSU CAFB, Ohio State University Companion Animal Fecal Bank; Lyo, lyophilized; Rep, replicate.

## 4. Discussion

The results of this study support our hypothesis that freshly processed canine and feline FMT products exhibit greater surrogate microbial viability compared to lyophilized FMT products. Our second hypothesis, that AnimalBiome^TM^ and CAFB lyo FMT products have comparable surrogate microbial viability, was partially supported and dependent on the species of origin. Canine CAFB Lyo products had significantly greater total CFU/g compared to DB and DBR FMT products; however, there was a high degree of overlap of the ranges of total CFU/g across these products. There was no statistically significant difference in total CFU/g between feline CAFB Lyo and KB FMT products. We also determined that microbial community structure varied by the donor of origin; and that FMT derived from a raw-fed dog (DBR) had significantly greater abundance of *Enterobacteriaceae* compared to two standard-fed canine fecal donors.

The FMT processing protocol for AnimalBiome^TM^ is proprietary; however, several similarities can be drawn between AnimalBiome^TM^ and CAFB FMT products. AnimalBiome^TM^ and CAFB fecal donors are rigorously screened to ensure donor health, which reduces the risk of disease transmission and possible transfer of detrimental phenotypes. Although outside the scope of this manuscript, rigorous and thorough FMT fecal donor screening is critical to ensure FMT product safety and consistency [33]. Another important similarity between AnimalBiome^TM^ and CAFB FMT is the use of glycerol as the preservative. Glycerol is listed as an ingredient in AnimalBiome^TM^ FMT products; however, the concentration is not disclosed. Glycerol was chosen as the cryopreservative in the CAFB FMT products evaluated herein to ensure as much similarity as possible between FMT products. CAFB FMT is mixed with glycerol to a final concentration of 10%, which is considered standard by many laboratories [54, 65, 66]. Though glycerol was used for all FMT products, the texture and consistency of AnimalBiome^TM^ and CAFB Lyo FMT products were very different (Figure 11). Despite 48 hours of lyophilization, CAFB Lyo FMT products were incompletely dry and sticky, which is consistent with findings in previous viability studies evaluating human-derived FMT products [43]. This sticky texture precludes homogenization to a fine powder and makes encapsulation subjectively difficult. In contrast, the AnimalBiome^TM^ FMT products were completely dry and homogenized. Small pieces of fur and undigested material were observed in AnimalBiome^TM^ FMT products, which were not observed in the filtered CAFB FMT products (Figure 11). This suggests that FMT processing, including filtration techniques and glycerol concentration, likely differ across these fecal repositories evaluated in this study.

**Figure 11.**
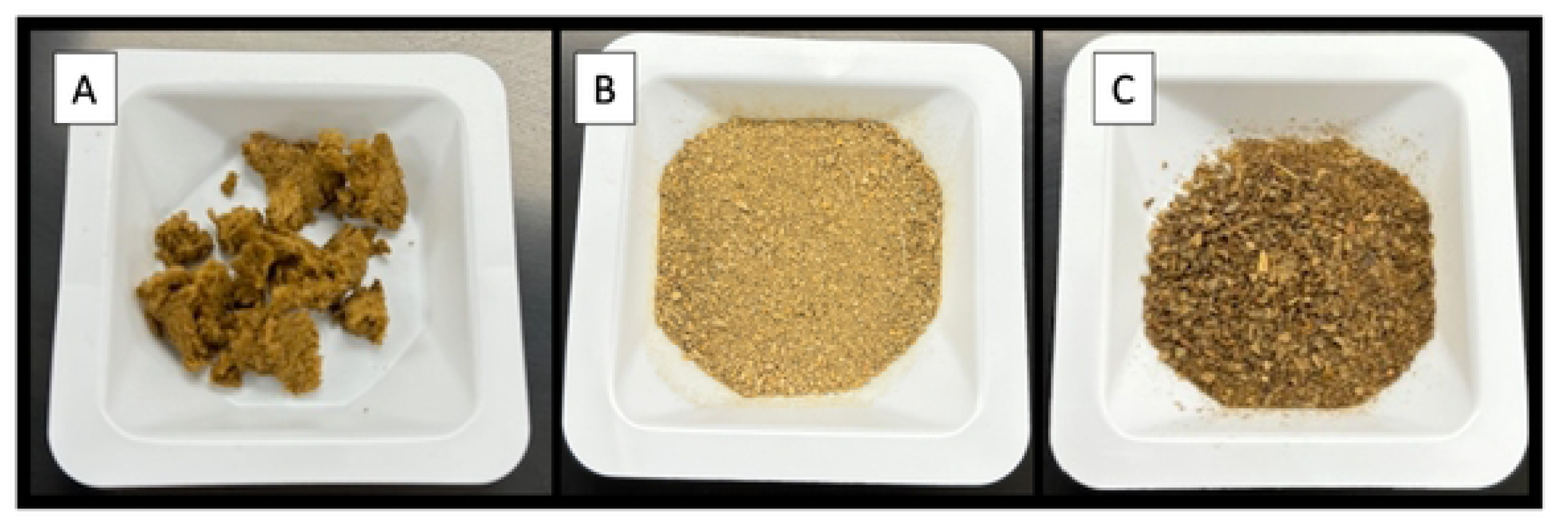
Comparison of texture and consistency across CAFB and AnimalBiomeTM products. A) CAFB Lyo product produced from a standard-fed canine fecal donor. The sticky texture precludes homogenization to a fine powder and makes encapsulation subjectively difficult. B) AnimalBiomeTM FMT from a standard-fed dog. The powder is completely dry and homogenized. C) AnimalBiomeTM from a raw-fed dog. The texture and consistency is similar to the standard-fed product. The texture of feline CAFB Lyo and KB products were similar to canine CAFB Lyo and AnimalBiomeTM products, respectively.

Lyophilization, or freeze-drying, is a popular method for preserving human and veterinary FMT. Lyophilization did not significantly impact the microbial community structure or alpha diversity of canine CAFB FMT products, indicating similar microbial compositions pre-and post-lyophilization. Of note, relative abundance is compositional and only indicates the presence of DNA and not the viability of the microbe. *In vitro* surrogate microbial viability testing, however, did show that lyophilization significantly decreases the total CFU/g in canine and feline CAFB FMT products. Gram-negative microbes were particularly impacted, with no detectable growth observed in the majority of lyophilized samples. These results are consistent with previous viability studies; which determined that the impact of lyophilization varies by microbe, characteristics of the suspension, and choice of cryopreservative [40, 67]. One study evaluating 10 microbial species showed that Gram-negative organisms tend to be more sensitive to lyophilization than Gram-positive organisms [67]. The cell wall structure of Gram-positive microbes likely plays a role in its resistance to lyophilization [68]. Human studies have shown that lyophilization is more detrimental to the viability and fitness of strict anaerobes than freezing; however, it yields a more shelf-stable product[40, 69].

Despite the loss of viability of certain microbes, lyophilized FMT is metabolically active [69] and capable of inducing remission of recurrent *Clostridioides difficile* in people [42, 44]. Further, two recent studies published by AnimalBiome^TM^ used 16S rRNA gene amplicon sequencing to evaluate microbial engraftment dynamics in 54 dogs and 46 cats with chronic gastrointestinal disease after a 25-day course of lyophilized oral FMT [70, 71]. In dogs, an average of 18% of the fecal donor’s bacterial ASVs engrafted in FMT recipients, with *Bacteroides* spp, *Fusobacterium* spp, and *Lachnoclostridium* spp engrafting more frequently than other taxa[70]. In cats, fecal donors shared an average of 13% of their bacterial ASVs with FMT recipients and the most commonly shared ASVs were *Prevotella 9*, *Peptoclostridium*, *Bacteroides*, and *Collinsella* [71]. Thus, despite having lower total CFU/g than fresh FMT products, lyophilized canine and feline FMT are biologically active and capable of shifting the microbial community structure of FMT recipients [70, 71]. These findings underscore the duality of lyophilization; while it reduces the viability of certain microbes, it preserves enough functionality to effectively modulate recipient microbial communities. This also highlights the need for further *in vivo* research to establish the ideal “dose” of viable microbes in order to optimally modulate the recipient’s microbiome and potentially improve clinical outcomes.

Alpha Diversity, which measures the microbial community diversity within individual samples (here an FMT product type), is a standard method for comparing microbial communities. In this study, DB FMT products exhibited significantly higher alpha diversity (Shannon Index and Observed ASVs) than all other canine FMT products. Diet is a well-established determinant of gut microbial composition and diversity. However, studies on the impact of raw versus kibble diets on canine fecal alpha diversity have yielded conflicting results [48, 51, 70]. One study of 27 raw fed dogs and 19 commercially fed dogs showed no significant difference in alpha diversity between diet groups[51]. Conversely, a study of eight healthy adult boxers found that Shannon Diversity increased after feeding a raw diet for 2 weeks [48]. Additionally, the previously mentioned study of 54 dogs published by AnimalBioime^TM^ found that raw diets were significantly associated with higher alpha diversity[70]. In the context of these previous studies, our results suggest that differences in alpha diversity observed between AnimalBiome^TM^ and CAFB FMT products may not be solely attributable to diet. Indeed, a plethora of other factors, including the FMT processing technique, patient body condition score, prior antibiotic administration, body size (large breed vs small breed), and lifestyle also contribute to microbial community structure[70]. Importantly, only one raw-fed donor and two standard-fed donors were evaluated in the study herein, making definitive comparisons between donors difficult. Other potential factors impacting alpha diversity in these products cannot be determined due to the proprietary nature of donor and processing information.

Studies in human medicine have highlighted the potential clinical importance of donor alpha diversity. For example, higher donor richness was associated with clinical success of FMT in people with refractory inflammatory bowel disease (IBD) [72]. A systematic review of 25 studies of people with ulcerative colitis found that higher alpha diversity of the FMT product was a predictor of clinical success [72]. Another metanalysis across various human FMT studies reported a significant positive association between alpha diversity and bacterial engraftment, but not clinical success [72]. There are a small number of studies that evaluate the impact of the donor microbiome on engraftment dynamics in dogs. The aforementioned study of 54 dogs published by AnimalBiome^TM^ used generalized linear models to identify host or donor factors that were associated with engraftment rates [70]. Their results indicate that ASV engraftment is not significantly associated with alpha diversity, but rather is impacted by the similarity between the donor’s and recipient’s microbiome pre-FMT [70]. These findings are similar to a study of 19 humans receiving FMT for recurrent *C. difficile infection*. Investigators used deep metagenomic sequencing to track strain-sharing between donor and recipients and determined that engraftment primarily depends on the abundance and phylogenic diversity of the donor and the recipient [73]. In canine medicine, the ideal level of alpha diversity required for effective therapeutic outcomes remains unknown. Other factors, such as specific donor-recipient pairings based on fecal microbial communities, may be a more important factor in determining engraftment. Ultimately, further research is needed to determine the impact of alpha diversity on microbial engraftment across multiple states of disease and donor-recipient pairs.

Beta diversity refers to the microbial diversity between different microbial communities, comparing how distinct or similar the microbial composition is between individuals. As expected, significant differences in beta diversity were observed between FMT products from different donors within each species regardless of FMT product type. This is consistent with findings from previous literature which show that individual dogs have unique microbial profiles analogous to a fingerprint[74–76]. Greater Bray-Curtis distances from the group centroid were observed within the DBR group compared to other canine FMT products (Figure 3B). This variation in microbial community structure was also reflected in a wider range of observed ASVs and Shannon Index in samples of DBR FMT. These findings may indicate that there is greater variability of microbial composition within DBR FMT products compared to others; however, a larger sample size that includes more than one donor per diet type is needed to draw more definitive conclusions.

Differential abundance testing was performed to compare microbial composition in standard fed and raw fed dogs. Previous studies have shown that the statistical method can significantly impact results and biological interpretations of microbiome data [77, 78]. Thus, for this study, a consensus of three differential abundance analyses was determined using LEfSe, MaAsLin2, and ANCOM-BC. A consensus of six differential abundant microbial families was established. Standard fed dogs (CAFB and DB) exhibit increased abundance of *Prevotellaceae* compared to the raw-fed dog. Five microbial families were more abundant in FMT products derived the raw fed fecal donor dog (DBR) including *Clostridiaceae, Enterococcaceae, Lactobacillaceae, Streptococcaceae,* and *Enterobacteriaceae.* One study of 27 raw fed dogs and 19 commercially fed dogs similarly identified increased abundance of *Enterococcus, Lactobacillaceae, Enterobacteriaceae,* and *Clostridium* by 16S rRNA gene amplicon sequencing; and increased *Streptococcaceae* detected by qPCR [51]. In contrast to the FMT products derived from the single raw-fed dog in this study, the raw-fed dogs in the previously referenced study exhibited a higher abundance of *Fusobacteria* and lower abundance of *Faecalibacterium* [51], which likely reflects differences in microbial community structures between individuals.

Increases in *Enterobacteriaceae* (*Escherichia* spp.) and *Streptococcaceae* (*Streptococcus* spp.) are associated with an abnormally increased dysbiosis index (DI), a fecal qPCR-based assay validated for use in dogs with chronic enteropathy [12]. However, this may not translate to deleterious effects on FMT recipients. A 2011 report from the American Academy of Microbiology emphasized that most strains of *E. coli* do not cause disease, and some are actually beneficial to the host [79]. For example, the *E. coli* strain Nissle 1917 is a probiotic that shows promise for improving gastrointestinal and urinary health in dogs [80] and is capable of adversely impacting *in vitro* growth of multidrug resistant uropathogenic *E. coli* in cats [81]. *Streptococcus salivarus* subsp. *thermophilus* is included in the multi-strain probiotic Slab51^®^. One study of 20 healthy dogs found that this probiotic is well-tolerated and capable of enhancing mucosal and systemic immune parameters [82]. A separate study of 20 dogs found that Slab51^®^ reduced clinical signs associated with colonic dysmotility and decreased inflammatory cell infiltrates [83]. In our study, *Escherichia* and *Streptococcus* were not characterized to the strain level, thus, it is unclear whether the enriched abundance in DBR products or viable colonies isolated from CAFB FMT products represent beneficial, detrimental or a combination of microbial strains.

Regardless, the Companion Animal FMT Consortium recommends the exclusion of canine and feline fecal donors if there is a history of pharmaceuticals or diets known to induce dysbiosis [33]. Specifically, the consumption of raw food within 30 days of screening precludes enrollment as a fecal donor [33]. Raw diets are often touted as natural, holistic, and healthy, however safety concerns exist and previous studies have demonstrated adverse alterations in the microbial composition of dogs [48, 49, 51, 52, 84–87] and cats fed a raw diet [85, 87, 88]; including decreased proportions of beneficial organisms such as *Lactobacillus* and *Faecalibacterium* and increased proportions of pathobionts, such as *Campylobacter jejuni*, *Escherichia coli, Salmonella*, and *Listeria monocytogenes* [48, 51, 52, 85]. The high prevalence of extended spectrum beta-lactamase producing microbes in the stool of raw-fed dogs is of particular concern, as it poses a risk for the spread of antimicrobial resistance genes [49, 51, 52, 86]. This not only endangers the health of the animal recipient, but also raises public health concerns due to the potential for zoonotic transmission of antimicrobial resistant pathogens to humans [89]. Despite these concerns, raw meat-based diets remain popular among owners and many veterinarians recommend them to their clients (author observations). Continued study of raw diets and its impact on the microbiome is warranted, particularly given evidence that the similarity of donor-recipient microbiomes is an important predictor of engraftment in dogs [70]. This may inform a personalized-medicine approach to match donor and recipient pairs based on their microbial compositions, which are largely dictated by diet.

Further, though differential abundance testing determined that FMT products derived from the raw-fed donor (DBR) had enriched abundances of ASVs classified as *Enterobacteriaceae, Streptococcaceae*, and *Clostridiaceae* compared to the standard fed dogs, only *Clostridiaceae* exhibited *in vitro* viability. In contrast, canine CAFB FMT products had lower abundances of these ASVs, but exhibited robust growth of *Streptococcaceae* and *Enterobacteriaceae*. This highlights the limitations of 16S amplicon sequencing in assessing microbial populations, as microbes detected may or may not be viable. Further, the detectable growth of microbial colonies *in vitro* may not translate to viability or functionality *in vivo;* and vice versa. Future clinical trials are needed to define the connection between viable microbes *in vitro* and their corresponding *in vivo* viability. Research evaluating the impact of lyophilization, storage temperature, and various cryopreservatives on surrogate microbial viability and microbial composition is ongoing; and will provide valuable insights for optimizing microbial diversity and viability in FMT products for companion animals.

In feline CAFB FMT products, *Bifidobacterium* was isolated from agar plates from fresh FMT products, but was not isolated in feline lyophilized FMT products. *Bifidobacteria* have long been considered beneficial to host health by aiding in fiber digestion and producing acetate [5]. *Bifidobacterim* spp. administered as single or multi-strain supplement is documented to improve stool consistency in dogs[90] and cats[91] and to reduce episodes of stress induced colitis in dogs in a kennel setting [90]. For these reasons, *Bifidobacteria* are commonly produced and lyophilized on industrial scales for probiotixc manufacture [92]. The loss of viability of this microbe may be due to osmotic stress, which is a major cause of cell damage incurred during lyophilization. The addition of sugar cryoprotectants are widely used to reduce osmotic stress and improve revivification potential of bacteria [43, 92, 93]. Non-reducing disaccharides, such as trehalose, have been shown to improve viability of freeze-dried *Bifidobacteria* spp and other probiotics [92]. Given these findings, it is postulated that trehalose may also increase overall viability and the diversity of viable microbes in lyophilized FMT products. In order to optimize microbial viability; and thus, the engraftment potential, of lyophilized FMT products, further research evaluating multiple cryopreservatives is needed.

Surrogate viable microbes isolated from canine CAFB FMT products were represented by three phyla: Firmicutes, Actinobacteria, Proteobacteria. In feline CAFB FMT products, an additional phylum, Bacteroidota, was identified. In contrast, surrogate viable microbes from both canine and feline AnimalBiome^TM^ FMT products were exclusively from the Firmicutes phylum. Separate culture experiments of lyophilized CAFB FMT products preserved with 10% glycerol show that Gram-negative bacteria are no longer viable at the one-month timepoint [94, 95]. Thus, the broader diversity of viable microbial phyla in CAFB Fresh and CAFB Lyo FMT products compared to AnimalBiome^TM^ FMT products is attributed to the temporal association of culture experiments relative to FMT processing. All CAFB FMT products were cultured immediately after processing (fresh) and again immediately after lyophilization was completed. Whereas, the time between the processing date and shipping date of AnimalBiome^TM^ FMT products is unknown. Though surrogate viable microbes in CAFB FMT products represent a more diverse group of microbial phyla, as mentioned above, lyophilized FMT products are effective, regardless if they are produced in academic laboratories or commercial settings [70, 71, 96]. Managing an in-house fecal bank requires substantial financial and labor commitments, which may be prohibitive for many veterinarians. Commercially available AnimalBiome^TM^ FMT products fill a critical niche in the veterinary industry by providing veterinarians and owners with consistent access to safe canine and feline FMT products.

## Limitations

This study evaluated the *in vitro* surrogate microbial viability of three commercially available canine and feline FMT products. We also compared the microbial composition in these FMT products, emphasizing comparisons between FMT derived from a raw-fed donor and two standard fed donors. Similar patterns of surrogate microbial viability were seen across FMT products and species and many findings correlate with previous work. However, there are some important limitations to this study. The first limitation is the small number of dogs and cats used for analysis (canine donors, n=3; feline donors, n=2). AnimalBiome^TM^ is a privately owned company that does not disclose detailed diet histories, lifestyle, or demographics of its fecal donors. We sought to maximize similarities between CAFB and AnimalBiome^TM^ FMT products by using the same cryopreservative; however, the processing technique, lyophilization dates and storage time of AnimalBiome^TM^ FMT products are not disclosed on package labels, making direct comparisons between CAFB Lyo FMT products and lyophilized AnimalBiome^TM^ FMT products difficult.

Microbial viability is often defined as the ability to produce progeny [97]; however, this is complicated by the fact that most intestinal microbes are not easily cultured using standard laboratory techniques [45, 46]. Thus, the total CFU/g yielded from each FMT product on selective agar represents only a fraction of potentially viable gut microbes, which herein are considered surrogates for the overall microbial community members within these FMT products. Further, 200 colonies were hand collected from each culture experiment and the presence or absence of these surrogate viable microbes are representative of all viable microbes grown on the agar plates. Finally, the metabolites produced by microbes were not evaluated, but are critically important for collaborative metabolism with host, cell signaling, and maintaining local and systemic health [3]. Future *in vivo* studies are needed to establish how processing techniques, cryopreservatives, and storage-time impacts the *in vitro* and *in vivo* metabolic potential of FMT in companion animals.

## Conclusions

Each individual fecal donor possesses a unique microbial profile. Six differentially abundance microbes were identified between raw fed and standard fed canine donors, which may have important implications when choosing fecal donor-recipient pairs. Fresh canine and feline FMT products exhibit robust growth of aerobic and anaerobic Gram-positive and Gram-negative microbes and are more representative of the original microbial assemblage of the fecal donor. Lyophilization of canine and feline FMT products, regardless of fecal repository, significantly decreases surrogate microbial viability, with particular impact on Gram-negative bacteria. While no specific therapeutic “dose” of viable microbes has been established, it is worth noting that 10^4^-10^7^ CFU/g within lyophilized FMT products represents a substantial number of viable microbes, similar to currently available probiotics [98]. Our study focuses on multiple lots of FMT, each produced from a single donor, which brings attention to the potential impact of donor-recipient pairing on engraftment success. Further studies are needed to evaluate the impact of donor-recipient pairs and “dose” of viable microbes on the *in vivo* growth and engraftment kinetics of fecal microbes to expand our understanding of the mechanisms underlying its therapeutic effects in companion animals.

## Acknowledgements

The authors would like to thank Dr. Nora Jean Nealon enrolling feline fecal donors in the Companion Animal Fecal Bank. The authors would also wish to thank the dedicated owners, as well as the dogs and cats enrolled in the fecal bank, for their time and commitment to this work.

